# Near-infrared phenomic and genomic prediction for seed protein in winter legume white lupin (*Lupinus albus* L.): A utility comparison

**DOI:** 10.64898/2026.08.05.743001

**Authors:** Mark Philip Castillo, Oluwaseye Gideon Oyebode, Ainsley Lenahan, Anthony Orloski, Marnin Wolfe

## Abstract

White lupin (*Lupinus albus* L.) is a cool-season grain legume with seed crude protein of 33-47%, competitive with soybean (*Glycine max* L.) meal. It also fixes nitrogen and mobilizes soil phosphorus. Because soybean is a summer crop, white lupin can occupy Southeastern winter fields as a complementary protein source. Breeding for seed protein is limited by the cost and throughput of reference phenotyping. To determine how each is best deployed, we compared the utility of near-infrared spectroscopy (NIRS)-based phenomic selection with genomic selection based on 246,847 SNPs from low-pass, whole genome sequencing in a panel of Auburn University breeding lines and USDA National Plant Germplasm System germplasm. A handheld NIR calibration against Dumas reference protein reached screening-grade accuracy (R^2^ = 0.81). Under common cross-validation, phenomic predictive ability was 0.93 and genomic was 0.12. The low genomic value was consistent with moderate heritability (H^2^ = 0.33) and strong genotype-by-year interaction. Beyond predictive ability, NIRS recovered superior accessions the strictest selection intensity, and 40 to 60 reference assays sufficed to calibrate the model. Handheld NIRS is a low-cost tool for protein calibration and early-generation screening, while genomic prediction remains suited to parental selection, together supporting a complementary strategy for legume breeding

**Plain Language Summary:** Soybean meal is the main protein source for livestock and fish farms in the United States. Because soybean is a summer crop, many Southeastern fields sit idle or grow low-value cover crops in winter. White lupin, a cool-season legume whose seeds are as protein-rich as soybean meal, makes a good complementary winter crop: it yields high-protein grain while serving as a cover crop that fixes nitrogen and frees up soil phosphorus for later crops. In our early-stage lupin breeding program, measuring seed protein by standard lab methods is slow and costly. We built a calibration that lets a handheld scanner estimate protein from light, and compared it with predicting protein from the plant’s DNA. The scanner gave accurate, low-cost protein screening from only about 40-60 lab tests, while DNA-based prediction remains suited to guiding parent selection. Used together, these tools offer breeders a practical path to develop high-protein white lupin.

**Core ideas:**

- Handheld NIRS provides screening-grade prediction of white lupin seed crude protein.
- Spectra carried more usable protein signal than markers by measuring seed chemistry directly.
- NIRS and genomic prediction serve different stages of a white lupin breeding program.
- About 40 to 60 reference assays sufficed to calibrate NIRS to near-full accuracy.

## Introduction

White lupin (WL) is a grain legume that combines high seed protein with soil-improvement services, making it a promising candidate for sustainable winter production in the Southeastern United States (Lucas et al., 2015; Quiñones et al., 2022; Noffsinger & van Santen, 2005). Mature seed typically contains about 33 to 47% crude protein (CP), competitive as an alternative feed protein source and above many cool-season legumes (Gresta et al., 2023; David et al., 2024). It fixes atmospheric nitrogen and mobilizes soil phosphorus through specialized cluster roots, improving nutrient cycling in nutrient-poor soils (Braum & Helmke, 1995; Quiñones et al., 2022). Also has been evaluated for adaptation in Alabama and the Southeastern United States (Noffsinger & van Santen, 2005).

Southeastern U.S. poultry and livestock production depends heavily on soybean meal and other feed ingredients transported into the region. In Alabama, animal agriculture consumed nearly 2.5 million tons of soybean meal in 202; yet the state’s soybean crop would supply only approximately 13% of that demand on a soybean-meal-equivalent basis (Caffarelli et al., 2025; Decision Innovation Solutions, 2022; USDA-NASS, 2022; U.S. Soybean Export Council, 2026). Meanwhile many winter fields remain fallow or grow cover crops that yield no marketable grain. WL offers a rare cool-season option, combining forage and cover-crop services with feed-grade seed protein (David et al., 2024; García-Gudiño et al., 2024; Lucas et al., 2015). Its feed value has been validated directly: WL supports broiler performance comparable to maize-soybean meal diets (Nalle et al., 2012) and can replace soybean meal in laying-hen and duck diets without major performance or health penalties (Laudadio & Tufarelli, 2011; Zapletal et al., 2017).

Among cool-season pulses, WL is distinguished by both high seed protein and slow genetic improvement. Faba bean (*Vicia faba*), field pea (*Pisum sativum*), lentil (*Lens culinaris*), and chickpea (*Cicer arietinum*) generally fall below its 33-47% protein range (Warsame et al., 2018; Hamungalu et al., 2026). Across these legumes, seed protein is quantitatively inherited and supported by comparatively recent genomic resources; pea and faba bean were until recently described as genomic orphans (Kaur et al., 2012). WL fits this profile, with genome-wide association studies for seed protein demonstrated only in the past few years (Schwertfirm et al., 2024; Annicchiarico et al., 2025).

Breeding programs need low-cost, rapid phenotyping, which remains a major bottleneck because laboratory assays are costly at breeding scale (Daba et al., 2022; Rife et al., 2021). Reference methods such as Kjeldahl and Dumas combustion provide reliable protein estimates but require chemical calibration and laborious laboratory workflows, making exhaustive accession-level testing impractical for large panels. (Mæhre et al., 2018; Daba et al., 2022). NIRS offers a rapid alternative by predicting seed composition from molecular absorbance associated with C-H, N-H, and O-H bonds after calibration to reference protein (Burns & Ciurczak, 2001; Daba et al., 2022). Portable NIRS further extends this approach to breeding programs needing inexpensive, high-throughput quality screening (Rife et al., 2021; Rincent et al., 2018). In parallel, genomic selection (GS) uses genome-wide markers to predict breeding values for genotyped but unphenotyped individuals, enabling earlier selection (Meuwissen et al., 2001; VanRaden, 2008). Recent studies in WL have demonstrated genome-enabled selection for grain protein, confirming protein as a quantitative but prediction-amenable breeding target (Schwertfirm et al., 2024; Annicchiarico et al., 2025).

These approaches converge in phenomic selection (PS), where NIRS or hyperspectral profiles are used like marker data to build relationship matrices for predicting quantitative traits (Rincent et al., 2018; Robert et al., 2022). In PS, spectra serve both as calibration inputs for chemical traits and as a measure of covariance among individuals, allowing BLUP-style models analogous to GS. Phenomic prediction has matched or exceeded genomic prediction in wheat (*Triticum aestivum*) and poplar (*Populus nigra*) (Rincent et al., 2018), wheat grain yield using hyperspectral relationship matrices (Krause et al., 2019), soybean (*Glycine max*) (Zhu et al., 2021), maize (*Zea mays*) (Lane et al., 2020), grapevine (*Vitis vinifera*) (Brault et al., 2022), and coffee (*Coffea canephora*) (Adunola et al., 2024). PS and GS are not interchangeable: GS targets transmissible additive genetic merit, whereas PS captures genetic, environmental, and tissue-specific variation expressed in the sampled material. The breeder’s question is therefore whether NIRS gives screening-grade protein estimates that can substitute for Dumas assays at breeding scale. Selection,including GS, depends on those phenotypes.

We evaluated Auburn breeding lines and USDA-NPGS accessions for seed crude protein across years using Dumas reference values, handheld NIRS spectra, and low-pass whole-genome SNP data. Our objectives were to: (i) develop and validate a handheld NIRS calibration for seed protein and estimate the heritability and genotype-by-year (GxY) interaction; (ii) compare phenomic and genomic prediction under a common cross-validation framework and test whether their integration improves prediction; and (iii) evaluate the operational value of NIRS prediction by measuring selection concordance across intensities for advancing high-protein and culling low-protein lines.

## Materials and Methods

### Plant material and field evaluation

The panel comprised WL accessions from two sources: advanced breeding lines from the Auburn University program, AU13L1082 cross series (70 accessions) and 134 USDA-NPGS accessions. The USDA-NPGS materials included 123 PI accessions and 11 accessions listed under other GRIN identifiers, including W and DLEG, and represented cultivars, breeding materials, and landraces of diverse origin. Trials were planted in the fall of 2023 and 2024, at the E.V. Smith Research Station - Plant Breeding Unit in Tallassee, Alabama, USA (32.5117° N, 85.8925° W). The experimental design in 2023 and 2024 was randomized completely block design (RCBD) with two replications, arranged in a 20 x 20 row-column grid. Each plot consisted of two 15-ft rows on raised ridges. After harvest, in order to ensure sufficient material for protein-determination, seed from replicate plots was combined within-year to generate year x genotype bulks for protein phenotyping and spectral scanning.

### Reference protein determination

Because Dumas analysis is resource-intensive, 150 accession-level seed bulks were selected, 30 (out of 187) from the 2023 and 120 (out of 193) from the 2024 field data, using a stratified diversity-sampling workflow described below. Consideration choosing a subset of 30 samples for 2023, is due to seed quality. Only 125 (2023 has 13 and 2024 has 112) out of 150 produced with usable spectra and genotype after preprocessing. Selection was stratified by plant architecture, alkaloid class, and seed size to preserve agronomic and end-use diversity, spanning determinate and indeterminate types, low- or no-alkaloid material, and small (0.5 cm), medium (1 cm), and large (1.25 cm). Within strata, samples were chosen to span the first principal component (explaining 62% of the variance) from a PCA on standardized total seed weight, dry biomass, plant height, and hundred-seed weight; these agronomic data were generated as part of a separate ongoing study and are not presented here (unpublished). Selected seed bulks were ground to 1 mm using a Retsch ZM 300 mill (RETSCH GmbH, Haan, Germany), and approximately 0.25 g of flour was combusted in tin foil cups on a LECO FP828P nitrogen analyzer (LECO Corp., St. Joseph, MI, USA) at the Auburn University Poultry Nutrition Laboratory. Each sample was analyzed in duplicate, with a third replicate run when duplicates differed by >0.15% N. Ethylenediaminetetraacetic acid (EDTA) calibration material (9.57% N), blanks, and reference materials were included daily, and accepted replicate values were averaged before conversion. Nitrogen was converted to as-received crude protein as CP (%) = N (%) x 6.25. Dry-matter content was determined gravimetrically on a parallel ~1.00 g sub-sample of each ground bulk, weighed into a pre-tared tin (standard tare 1.265 g tin foil cups; total initial mass ≈ 2.265 g), oven-dried to constant mass at 105 °C in 14 hours, and re-weighed using the formula DM (%) = [(W_final - W_tin) / 1.0] x 100, where W_final is the post-drying mass of tin plus dried seed (g) and W_tin is the tin tare (g). Crude protein was then expressed on a dry-matter basis as CP_DM (%) = CP_as-received (%) / [DM (%) / 100].

### NIRS spectral acquisition and preprocessing

Spectra were acquired from ground seed flour of 187 (2023) and 189 (2024), a total of 376 samples using a handheld MEMS-based FT-NIR spectrometer (NeoSpectra, Si-Ware Systems, Menlo Park, CA, USA). Each sample was scanned 13 seconds (maximum range) with reshaking between scans to capture packing and particle-size heterogeneity, and the 5 technical spectra replicates were averaged to one raw spectrum per accession. Spectra were harmonized to a common wavenumber grid and linked to accession, year, and CP source using dplyr, and data.table in R (R Core Team, 2024; Wickham et al., 2026; Barrett et al., 2026). Raw absorbance covered approximately 1350 to 2550 nm and was trimmed to the informative 1,389-2,222 nm region, then preprocessed by standard normal variate correction (Barnes et al., 1989) and Savitzky-Golay second-derivative transformation (polynomial order 2, window 11, derivative 2; Savitzky & Golay, 1964) in prospectr v0.2.10 (Stevens & Ramirez-Lopez, 2026), yielding 188 out of 257 spectral predictors (Figure 1b).

**Figure 1.**
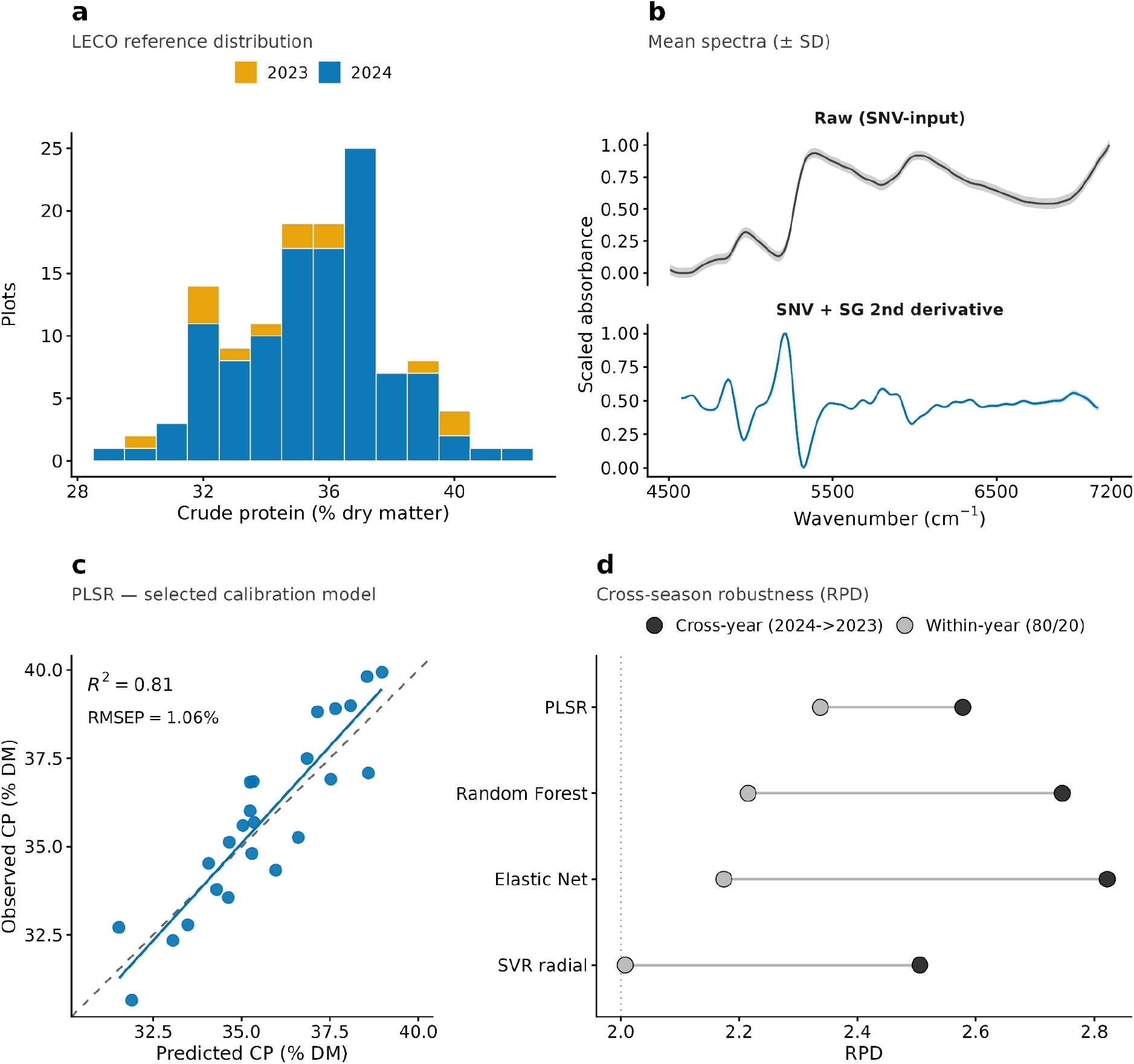
Handheld NIRS calibration for seed crude protein in white lupin. (a) LECO Dumas reference protein across calibration plots by season (2023, 2024), spanning 28.96 to 42.11% dry matter. (b) Mean seed spectra before and after preprocessing: raw scaled absorbance (top) and the standard normal variate (SNV) plus Savitzky-Golay second-derivative transform (bottom; polynomial 2, window 11) over the retained 1,389-2,222 nm region (mean ± SD). (c) Observed versus predicted protein for the selected PLSR calibration under 80/20 validation (R^2^ = 0.81, r = 0.91, RMSEP = 1.06%); solid line, linear fit; dashed line, 1:1. (d) Cross-season robustness of four calibration models, expressed as the ratio of performance to deviation (RPD; the standard deviation of the reference values divided by RMSEP, where higher values indicate better prediction relative to the trait’s natural variation). Two validation schemes are compared: within-year (80/20; grey) and cross-year transfer (train 2024, predict 2023; black), which respectively assess prediction accuracy within a season and transferability to an unseen season. All models exceeded the screening-grade threshold (dotted line, RPD = 2; Williams, 2014) under both schemes, indicating the calibration retains screening-grade accuracy when transferred to an unseen season.

### NIRS calibration and validation

Calibrations between preprocessed spectra and reference CP were developed at the accession-year level. Partial least squares regression (PLSR) was used as the primary model because it is well suited to high-dimensional, collinear spectral predictors (Wold et al., 2001). The number of latent components was optimized by cross-validation, and PLSR was benchmarked against elastic net (Zou & Hastie, 2005), random forest (Breiman, 2001), and radial-kernel support vector regression (Drucker et al., 1997) under identical preprocessing. Performance was evaluated using 80/20 cross-validation and cross-year validation between 2023 and 2024, with accuracy summarized by the coefficient of determination (R^2^), the root mean squared error of prediction (RMSEP), the ratio of performance to deviation (RPD = SD_reference / RMSEP), the mean absolute error (MAE) (Williams, 2014; Figure 1c,d). The selected PLSR model was refitted using all paired samples and applied across the panel. NIRS preprocessing and calibration were performed locally in R.

### Quantitative-genetic analysis

Variance components for of each accession CP were estimated by REML using lme4 (Bates et al., 2015):

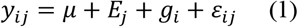

where y_ij_ is the CP of accession i in year j, μ is the overall mean, E_j_ the fixed year effect, g_i_ ~ N(0, σ^2^g) is the random accession effect, and ε_ij_ ~ N(0, σ^2^e) the residual.

Broad-sense heritability on a line-mean basis was estimated by the Falconer formulation

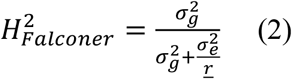

where 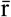 is the harmonic mean number of observations per accession (Holland et al., 2003). For the unbalanced design, Cullis heritability was also calculated as;

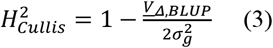

where 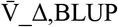 is the mean variance of pairwise differences among accession BLUPs (Cullis et al., 2006; Piepho & Möhring, 2007). Genetic coefficient of variation was computed as;

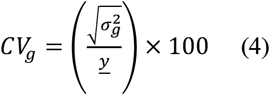

Genotype by year stability was assessed using the Pearson correlation of CP between years. Accession BLUPs were extracted as the primary prediction target (i.e. the data predictions are validated against), while BLUEs from a parallel fixed-accession model were retained for analyses requiring unshrunk accession estimates.

### NIRS statistical validation

NIRS calibration accuracy was quantified on independent validation sets using five metrics: R^2^, RMSEP, RPD, MAE, and the mean bias:

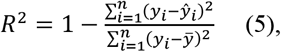

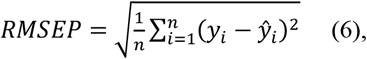

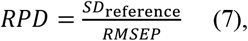

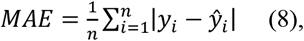

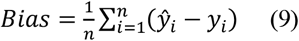

where y_i_ and ŷ_i_ are observed and predicted protein and 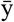 is the mean of observed values. An RPD ≥ 2.0 is adequate for screening and ≥ 2.5 for quality classification (Williams, 2014).

### Calibration efficiency

To determine how many Dumas assays were needed to calibrate the NIRS model, we built a learning curve over calibration-set size. For each size n_d ∈ {10, 15, 20, 30, 40, 60, 80, 95} we drew 200 random subsamples without replacement (Monte Carlo cross-validation; Picard and Cook, 1984). Each reserved a fixed 30-sample test set and drew n_d training samples from the remaining records, disjoint from the test set; the constant test size keeps accuracy comparable across sizes. Spectra were preprocessed by standard normal variate transformation and a second-derivative Savitzky-Golay filter in prospectr, and a PLSR model was fitted per subsample with pls, with latent components chosen within each training set by cross-validation under the one-standard-error rule (maximum 15). Each subsample gave the accuracy r (Pearson correlation of predicted and measured protein on the held-out set), the RMSEP, and the entry-mean heritability of the predicted protein, from CP ~ Year + (1 | Accession) fitted by REML in lme4 (Bates et al. 2015) as 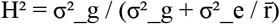 (Eq. 2), where σ^2^_g and σ^2^_e are the among-accession (genetic) and residual variances and 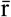 is the harmonic-mean number of records per accession. For each size we estimate the mean and 2.5th to 97.5th percentile (Figure 4) interval across subsamples, each metric scaled to its value at the largest training set (n = 95), the maximum allowed after reserving 30 of the 125 spectra-matched dumas assay of records for testing.

### Genotyping and relationship matrices

Single nucleotide polymorphisms (SNPs) were generated by low-pass whole-genome sequencing of the 354-accession WL diversity panel using the KHUFU pipeline (https://www.hudsonalpha.org/khufudata/; Korani et al., 2021), aligned to WL reference genome (Amiga variety, https://www.whitelupin.fr/; Hufnagel et al., 2020). Low-pass whole genome sequencing (WGS) yielded 256,652 cleaned SNP calls across the 354 accessions; after filtering for minor allele frequency (MAF) ≥ 0.01 using PLINK v1.9 (Chang et al 2015) across the full panel, 247,836 biallelic SNPs remained. Of the 354 accessions, the 204 with matching protein phenotypes, spectra, and were retained for prediction; markers that were monomorphic within this subset were removed, leaving 246,847 biallelic SNPs for these 204 accessions. The genomic relationship matrix (GRM) was computed by VanRaden’s (2008) first method, G = ZZ^T^ / [2Σ_j_p_j_(1−p_j_)], with Z the centered dosage matrix, and the phenomic relationship matrix (PRM) was built analogously from accession-mean preprocessed spectra (Eq 10). Both matrices were scaled to a unit mean diagonal for variance-component comparability (Forni et al., 2011; Technow et al., 2013; Figure 2a,b). This SNP density exceeds that of earlier WL studies based on reduced-representation genotyping such as GBS, SeqSNP, or DArT-seq (Schwertfirm et al., 2024; Surma et al., 2025; Annicchiarico et al., 2025).

**Figure 2.**
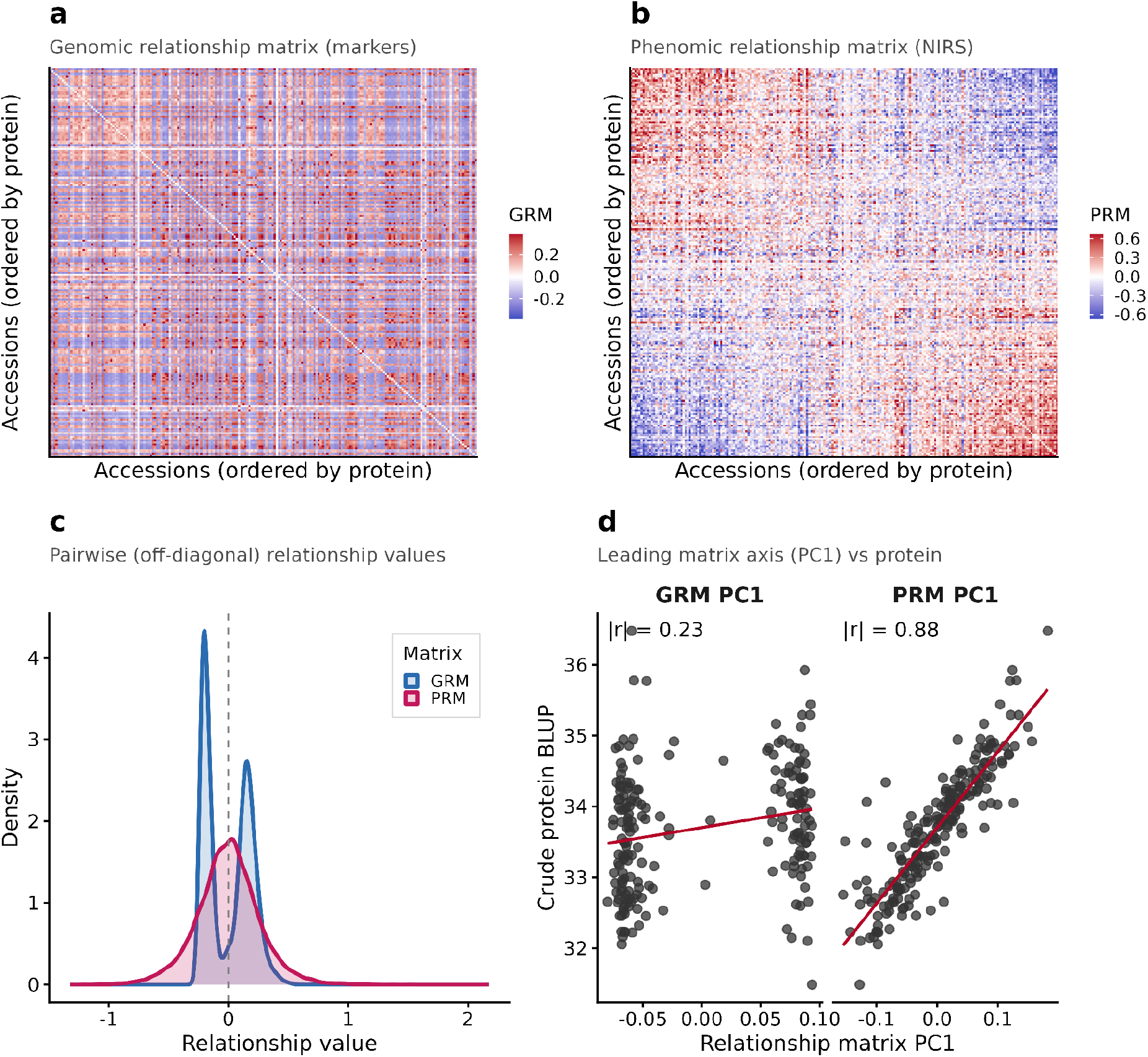
Genomic and phenomic relationship matrices and their alignment with seed crude protein. (a) Marker-derived GRM among the 204 common accessions, ordered by protein with the diagonal masked to emphasize off-diagonal structure. (b) NIRS-derived PRM, scaled to mean diagonal one and ordered identically. (c) Off-diagonal pairwise relationship distributions for the GRM and PRM, showing distinct structural profiles. (d) Association between each matrix’s leading axis (PC1) and protein, reported by magnitude since PC sign is arbitrary. The genomic axis was weakly associated (|r| = 0.23) and the phenomic axis strongly associated (|r| = 0.88).

PRM was built analogously from accession-mean preprocessed spectra. With S the n x k spectral matrix (k = 188 bands after preprocessing) and Sc its column-centered form

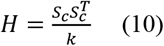

Each matrix K ∈ {G, H} was scaled to a mean diagonal of one for comparable variance components (Forni et al., 2011; Technow et al., 2013)

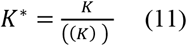

We also tested Gaussian kernels, computed from squared Euclidean distances D^2^ between standardized profiles,

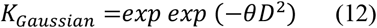

with bandwidth θ = 1 / median(D^2^); all kernels were diagonal scaled (Eq. 11) before fitting.

### Genomic and phenomic prediction

Genomic and phenomic predictions were compared in a common two-stage mixed-model framework. In the first stage, accession CP values were obtained from best linear unbiased predictors (BLUPs) from the quantitative-genetic model (Eq. 1), fitted from the dataset of 125 Dumas-assayed records and 251 NIRS-predicted records. Best linear unbiased estimators and (BLUEs) were also estimated for comparison (Supplementary table S1). In the second stage, these values were predicted using either genomic or phenomic predictors, using several different prediction models (see below).

Two cross-validation schemes were applied: repeated five-fold cross-validation (50 repeats) and repeated 80/20 validation (100 repeats). Within each repeat, a single set of train-test partitions was generated and applied to every model using both genomic and phenomic predictors, so all comparisons were paired across identical test sets. In each cycle, test-accession values were masked, the model was trained on the remaining accessions, and masked values were predicted from the corresponding predictors. Predictive ability (PA) was measured as the Pearson correlation between predicted and observed test values:

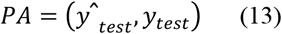

where ŷ_test is the vector of predicted CP values for the masked accessions and y_test is the corresponding observed vector. Genomic and phenomic predictors were then integrated using a two-kernel GBLUP model, in BGLR (Pérez & de los Campos, 2014):

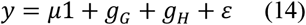

where g_G and g_H are the random genomic and phenomic effects modeled through K = G and K = H, respectively, and the remaining terms are as defined above.

### Prediction models

We compared three types of prediction models: kernel-based mixed models, Bayesian regressions, and machine-learning regressions. We used mixed.solve in the rrBLUP v4.6.3 package (Endelman, 2011) to fit a single-kernel GBLUP (using the Van Raden matrix) and RKHS (using the gaussian kernel matrix). We used the BGLR v1.0.9 package to fit two Bayesian regression models, BayesA and Bayes B (Meuwissen et al., 2001; Pérez & de los Campos, 2014). We fit a partial least squares regression (PLSR; Wold et al., 2001), a linear dimension-reduction method, using pls v2.9 package (Liland et al., 2026). Finally, two non-linear models, random forest with 500 trees (Breiman, 2001) was fitted using using randomForest v4.7-1 (Liaw & Wiener, 2002), and support vector regression with radial and linear kernels (Drucker et al., 1997) using e1071 v1.7-17 were tested. All seven models were evaluated under the cross-validation schemes as described above. Bayesian models in the model comparison used 6,000 iterations with a 1,000-iteration burn-in and thinning of five.

Multi-source integration was tested with a two-kernel RKHS model, y = μ1 + g_G + g_H + ε (BGLR; 8,000 iterations, 2,000 burn-in, thinning five) using 20 repeats of five-fold cross-validation, to compare single-and two-kernel models (Supplementary Table S3). Because Gaussian-kernel accuracy can depend on the bandwidth (Pérez-Elizalde et al., 2015), we check the kernel at 0.25θ_0_, 0.5θ_0_, θ_0_, 2θ_0_, and 5θ_0_, where θ_0_ is the baseline defined in Eq. 12, under the same cross-validation scheme. Predictive ability was insensitive to this choice: genomic accuracy was stable across all five bandwidths, and phenomic accuracy was stable from 0.25θ_0_ to θ_0_ and declined only at wider settings. We therefore kept θ_0_ at its original value for all Gaussian-kernel results. The linear kernel outperformed the Gaussian throughout, so all headline results use the linear kernel. Within group cross-validation was performed within the AU and NPGS groups under five-fold cross-validation, 50 repeats (Supplementary Table S4), and cross-year prediction ability was evaluated for all models in both directions (Supplementary Table S5).

### Selection concordance

Because a prediction method is useful only if it selects the same superior lines as direct measurement, we compared accessions selected by cross validated kernel prediction of CP with those selected by observed CP. Predicted CP was the cross-validated kernel-BLUP prediction from the GRM or PRM. At each selection intensity π, we retained the top k = ⌈πn⌉ accessions ranked by predicted CP (S_pred) and by observed CP (S_obs), where n is the number of accessions rank;the same procedure on the bottom k evaluated negative selection (culling of low-protein lines). Overlap was summarized as:

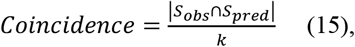

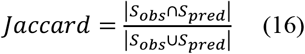

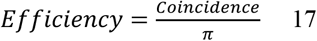

where coincidence is the fraction of superior accessions recovered by prediction, the Jaccard index is the overlap relative to the union of both selections, and efficiency is the enrichment over random selection (1 = chance). Concordance was evaluated at π = 0.10, 0.20, and 0.30 using the same five-fold, 50 repeated cross-validation used for the model comparison, with genomic and phenomic predictions generated on identical fold assignments within each repeat. Overall rank agreement was summarized by Spearman’s correlation. Quantitative-genetic, genomic, and phenomic prediction analyses were run in R 4.4.1 (R Core Team, 2024) on the Auburn University Easley High Performance Computing cluster (AU-HPC), using compute nodes with 43 cores and 350 GB memory under interactive and SLURM scheduler modes.

## Results

### Handheld NIRS calibrates seed protein accurately and transfers across seasons

Handheld NIRS calibration against Dumas reference protein supported downstream analyses. Across 125 accession NIRS-LECO samples spanning 28.96-42.11% protein, PLSR predicted crude protein with R^2^ = 0.81, RMSEP = 1.06%, RPD = 2.34, and bias = −0.16%, MAE = 0.97 under repeated within-panel 80/20 hold-out validation (Figure 1c; Supplementary Table S2). PLSR was retained because random forest, elastic net, and radial SVR did not improve performance (RPD = 2.22, 2.17, and 2.01, respectively; Figure 1d).

### Seed protein is moderately heritable

Heritability for CP was H^2^ = 0.34 using the Cullis formulation and H^2^ = 0.33 using the line-mean variance-component formulation, with genotypic and residual variances of 1.56 and 4.73, respectively. CP was year-dependent, increasing from 33.5% in 2023 to 35.2% in 2024. The cross-year phenotypic correlation of accession means was low on both raw means (r = 0.28; Figure 3a).

**Figure 3.**
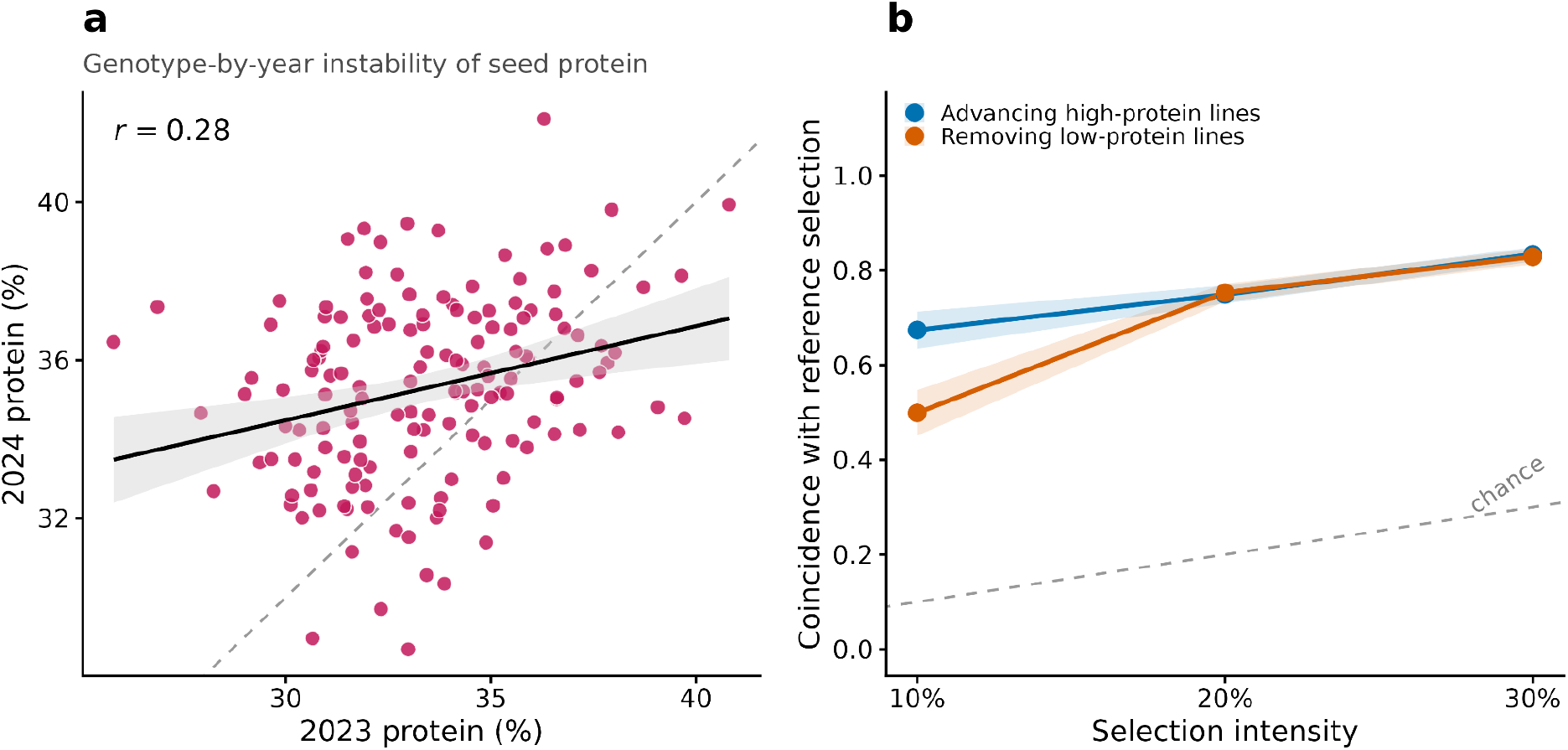
Phenomic and genomic prediction of seed crude protein, and the quantitative-genetic limits on predictability. (a) Accession-mean protein in 2023 versus 2024 (n = 125 accessions in both seasons); solid line and band, linear fit with 95% CI; dashed line, 1:1. The weak cross-year raw phenotypic correlation (r = 0.28) indicates substantial genotype-by-year interaction. (b) Selection concordance between predicted and reference-selected accessions at matched selection intensities (n = 204; 50 cross-validation cycles). Bands, ±1 SD; dashed line, chance. Advancing outperformed removal at the strictest intensity and the two converged as selection relaxed.

### Phenomic prediction and genomic prediction performance

We evaluated genomic and phenomic prediction on the same 204 accessions, the same protein target, and the same prediction model. The only difference was the relationship matrix / predictor set. Under five-fold cross-validation, phenomic prediction achieved higher predictive ability than genomic prediction (0.93 versus 0.12), and this persisted under repeated within-panel 80/20 hold-out validation of genomic prediction at 0.19 and phenomic prediction at 0.89. The leading axis of the PRM was strongly associated with protein (|r| = 0.88), whereas the leading genomic axis was weakly associated (|r| = 0.23; Figure 2d).

### The phenomic advantage is robust and not improved by genomic integration

The phenomic prediction ability was robust across model classes and kernel choices. Across seven prediction models, marker-based predictive ability remained low, whereas spectra-based predictive ability was consistently higher (Table 1). Gaussian kernels gave no improvement over linear kernels for either relationship matrix, and genomic prediction was insensitive to the Gaussian bandwidth. Combining both matrices in a two-kernel model gave no improvement over the phenomic kernel alone when both were fitted under the same solver and identical partitions (0.892 versus 0.894; Δ = 0.002; Supplementary Table S3).

**Table 1.**
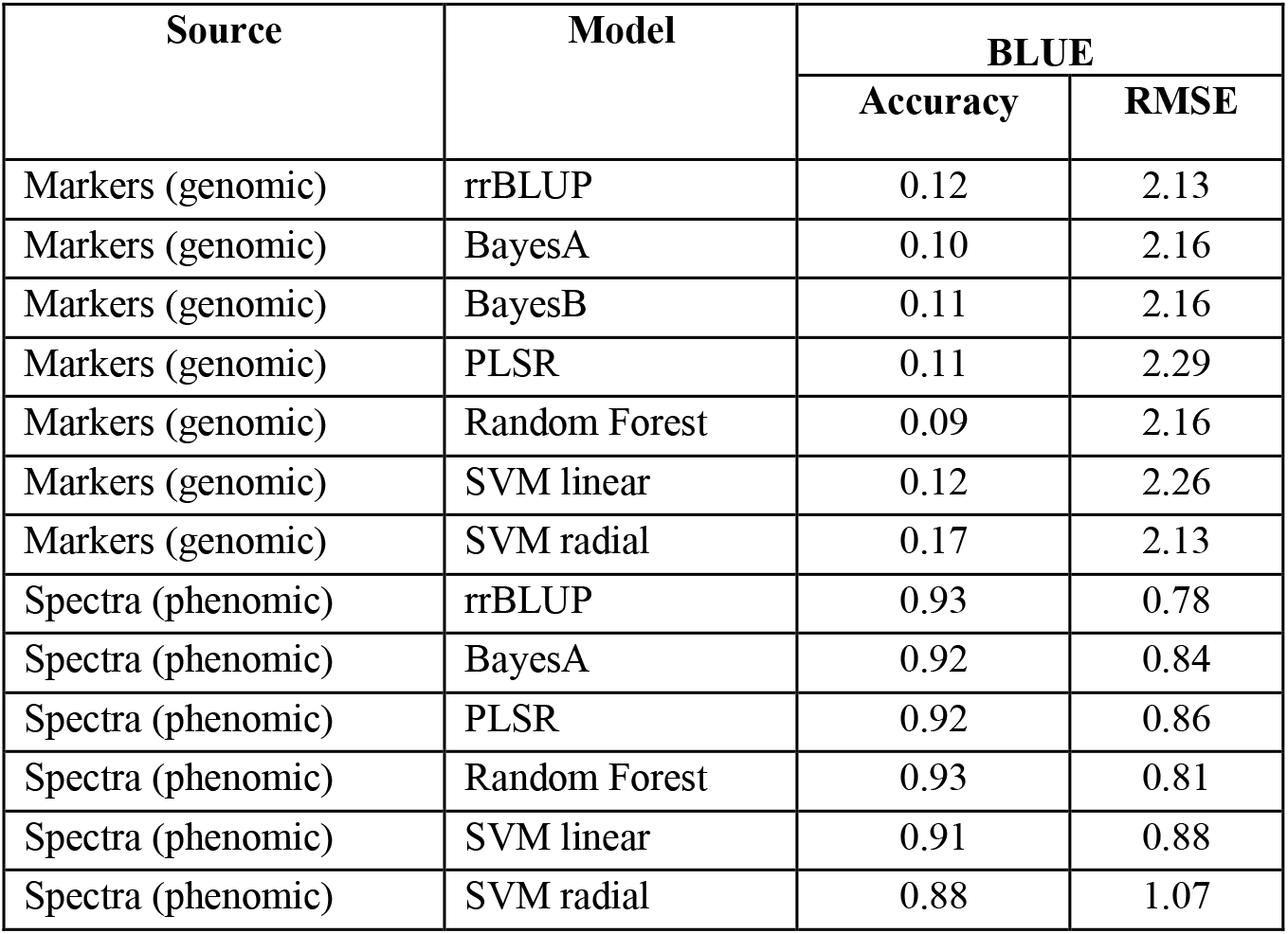
Predictive ability of seven genomic and phenomic prediction models. Predictive ability (Pearson correlation between predicted and observed values) and RMSE for genomic (markers) and phenomic (spectra) prediction under 50 replicates of five-fold cross-validation on the same 204 accessions, using the accession BLUE as the target.

| Source | Model | BLUE |  |
| --- | --- | --- | --- |
|  |  | Accuracy | RMSE |
| Markers (genomic) | rrBLUP | 0.12 | 2.13 |
| Markers (genomic) | BayesA | 0.10 | 2.16 |
| Markers (genomic) | BayesB | 0.11 | 2.16 |
| Markers (genomic) | PLSR | 0.11 | 2.29 |
| Markers (genomic) | Random Forest | 0.09 | 2.16 |
| Markers (genomic) | SVM linear | 0.12 | 2.26 |
| Markers (genomic) | SVM radial | 0.17 | 2.13 |
| Spectra (phenomic) | rrBLUP | 0.93 | 0.78 |
| Spectra (phenomic) | BayesA | 0.92 | 0.84 |
| Spectra (phenomic) | PLSR | 0.92 | 0.86 |
| Spectra (phenomic) | Random Forest | 0.93 | 0.81 |
| Spectra (phenomic) | SVM linear | 0.91 | 0.88 |
| Spectra (phenomic) | SVM radial | 0.88 | 1.07 |

### Genomic prediction does not transfer across seasons

Cross-year predictions were conducted wherein one season was used to predict the other. Accuracy was consistently low across all seven models in both directions (forward 2023 to 2024, 0.14; backward 2024 to 2023, 0.12; Supplementary Table S5); the cross-year correlation was 0.31. Genomic predictive ability plateaued near 0.2 by approximately 100 training accessions and did not increase with larger training sets (Supplementary Table S7). Genomic prediction was positive in the pooled panel but negative within each germplasm source (AU = −0.23; NPGS = −0.18; (Supplementary Table S4).

### NIRS prediction for selection

Selection concordance was evaluated in two ways. First, across 50 cross-validation cycles, NIRS-predicted protein recovered 47%, 69%, and 79% of the reference-selected high-protein accessions at the 10%, 20%, and 30% selection intensities, corresponding to 4.7, 3.4, and 2.6 times the recovery expected by chance (Spearman r = 0.91; Figure 3b;Supplementary Table S8). Second, comparing advancement against culling at the strictest intensity, prediction recovered 67% of the truly high-protein accessions but 50% of the truly low-protein ones (6.7 and 5.0 times chance); the two directions converged at 20% and were indistinguishable at 30% (Figure 4b). Calibration efficiency plateaued near 40 reference assays. At 40 assays, calibration accuracy reached 98% of its full-set value (r = 0.89), RMSEP came within 8% of the full calibration (1.18 vs 1.09% DM), and the heritability in the predicted values reached 98% of full (Figure 4).

## Discussion

To our knowledge, this is the first direct comparison of phenomic and genomic prediction for seed CP in white lupin. It extends phenomic prediction to a high-protein cool-season legume, after prior work concentrated on cereals, oilseeds, and woody or perennial species (Rincent et al., 2018; Robert et al., 2022; Zhu et al., 2021; Brault et al., 2022; Adunola et al., 2024; DeSalvio et al., 2024).

We report three practical findings. (1) Handheld NIRS calibrated seed protein at screening-grade accuracy and transferred across years, requiring 40 to 60 reference assays to build. (2) Phenomic prediction reproduced protein far more accurately than genomic prediction (0.93 and 0.12), and combining the two gave no further gain. Phenomic prediction estimates the phenotype while genomic prediction the breeding value, so the two accuracies are not the same quantity (Feldmann et al., 2026; Wang et al., 2025). (3) At the strictest intensity, prediction recovered high-protein lines more reliably than it removed low-protein ones (0.67 vs 0.50); at moderate intensity the two were comparable. Because a strict cut at screening grade risks discarding useful germplasm, prediction can be applied at moderate intensity (top 30%), where 79% of reference-selected lines are recovered.

### A low-cost calibration lowers the reference-phenotyping barrier

The calibration addresses a major bottleneck in protein-oriented breeding: the cost and throughput of phenotyping. Dumas combustion is reliable but difficult to scale, whereas handheld NIRS delivered screening-grade estimates from a ground-seed scan performed in-house, and those estimates transferred across years. This allows more lines to be screened per cycle in early-stage and resource-limited programs. Consistent with the practical appeal of phenomic approaches, as low-cost and high-throughput alternatives to marker-heavy workflows (Rincent et al., 2018; Lane et al., 2020; Weiß et al., 2022). Our calibration accuracy (R^2^ = 0.81, RMSEP = 1.06%) matched or exceeded published legume NIRS protein calibrations, including WL work on chemically referenced samples and pea protein models reporting R^2^ of 0.72 to 0.79 (Daba et al., 2022), and did so with a low-cost handheld instrument rather than a benchtop system. Calibration was also efficient. Accuracy, error, and the heritability in the NIRS values plateaued around 40 assays, reaching about 98% of their full-set values, with little gain thereafter (Figure 4). At 40 the average calibration is already strong, but performance still varies with which samples are used; by 60 that variation is small, so any single calibration of this size performs well. A reference set of 60 determinations therefore suffices both to calibrate the instrument within our population, and to recover the genetic signal, after which further wet chemistry adds little.

**Figure 4.**
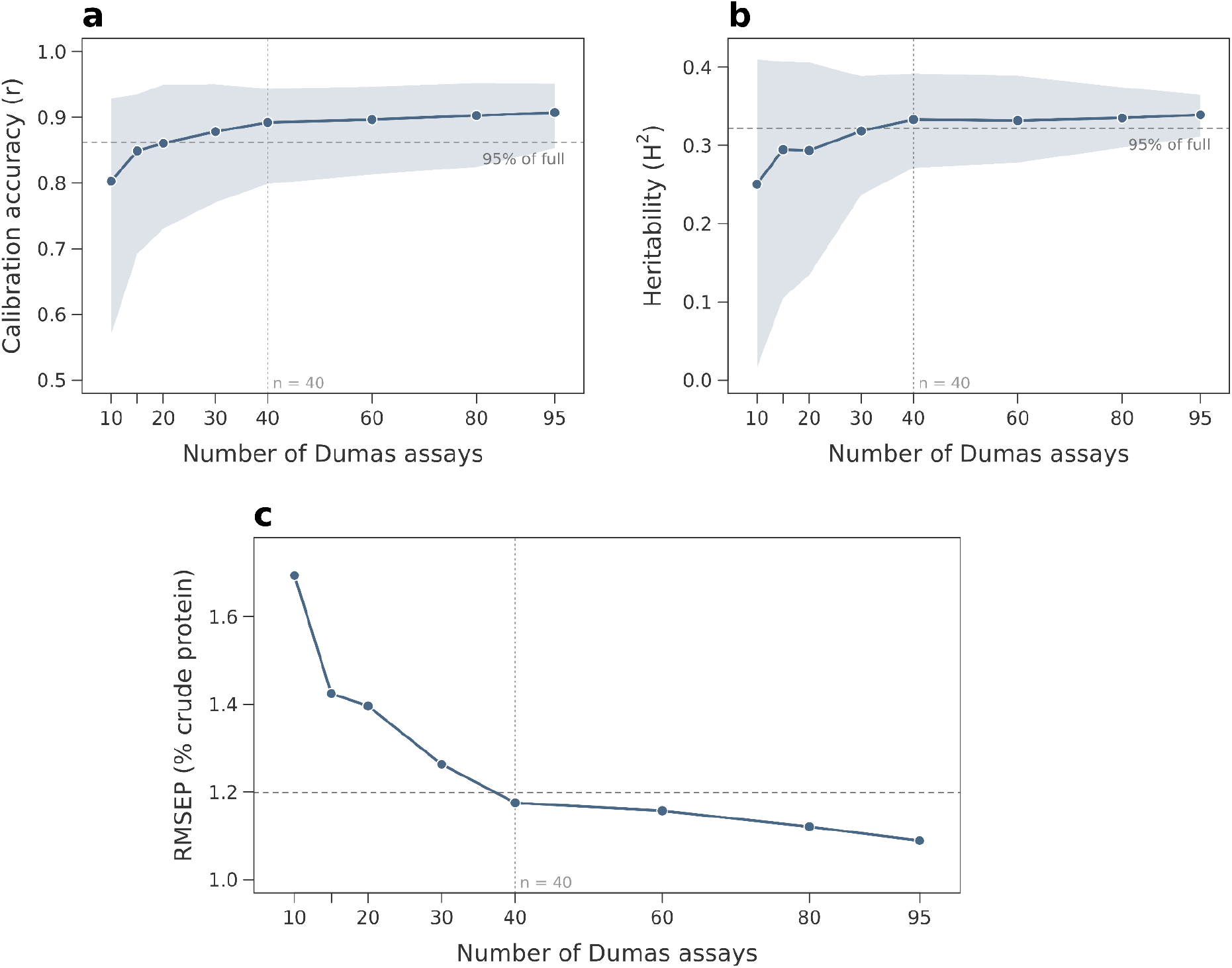
NIRS calibration efficiency: a modest reference set reaches near-full calibration. Calibration accuracy (Pearson r, panel a), entry-mean heritability in NIRS-predicted protein (R^2^, panel b), and RMSEP (panel c) as the PLSR calibration set increases from 10 to 95 Dumas-assayed records. Points are means over 200 random subsamples (Monte Carlo cross-validation), each holding out a fixed 30-record test set; shaded bands are the 2.5th to 97.5th percentile ranges. Dashed lines mark 95% of the full-calibration value (panels a, b) and 10% above the full-calibration RMSEP (panel c); the vertical guide marks n = 40. All three metrics stabilized near 40 records, where accuracy reached 98% of full (r = 0.89) and RMSEP came within 10% of its full value (1.18 vs 1.09% crude protein).

### Why spectra and markers capture different signals

PS exceeded genomic prediction for seed CP under identical validation (0.93 and 0.12), consistent with studies where NIRS-or hyperspectral-derived relationships matched or surpassed genomic relationships for complex traits (Rincent et al., 2018; Krause et al., 2019; Zhu et al., 2021; WeiB et al., 2022; Adunola et al., 2024). The likely explanation is that the two matrices capture different biological signals: the GRM reflects variation in genomic relatedness as captured by markers, whereas the PRM captures realized seed biochemical composition, i.e. phenotype-level variation, which combines genetic and environment-expressed variation in the sampled seed. Spectra therefore behaved as phenotypic-proxy, supported by the leading-axis analysis where the PRM axis tracked protein more strongly than the GRM axis (|r| = 0.88 vs. 0.23; Figure 2d). Because spectra encode direct chemical information on protein, phenomic predictive ability partly reflects measurement rather than prediction of independent genetic merit and is not strictly comparable to genomic accuracy (Wang et al., 2025).

This difference should not be described as phenomic prediction superiority. The phenomic result was consistent across all models, including non-linear machine-learning regressions. Gaussian kernels did not improve genomic prediction, and within-group GS collapsed to negative values. Likewise, combining the genomic and phenomic kernels gave no gain over the phenomic kernel alone when both were fitted under the same solver (Δ = 0.002). Indicating that adding the genomic kernel did not improve predictive accuracy. Because phenomic features are downstream of the same genotype and environment as the target, the two kernels are not independent, so this model is used for prediction only, not to partition signals (Feldmann et al., 2026). Genomic prediction remains suited to parental choice and ranking unscanned candidates, because breeding values estimate transmissible genetic merit and these uses precede harvested seed. Thus, PS is best positioned as a rapid upstream screen for within-season screening of harvested seed, while GS remains a downstream tool for crossing and long-term improvement and serves parent selection as related training material, environment-specific training sets accumulate (Meuwissen et al., 2001; VanRaden, 2008; Bernardo, 2020; Gebremedhin et al., 2023; Schwertfirm et al., 2024; Annicchiarico et al., 2025; Feldmann et al., 2026).

### What predictive ability means for selection

Predictive ability is a statistical measure. Its breeding relevance depends on whether predictions recover the right genotypes at practical selection intensities. At matched intensities in selection concordance (Figure 3b), phenomic prediction recovered two-thirds of the superior accessions at the strictest selection, 6.7 times chance. High predictive ability reflects agreement with the Dumas phenotype, not realized genetic gain, which additionally depends on the heritability of the spectral index and its genetic correlation with the trait (Feldmann et al., 2026). Genomic prediction was weak, the predictive ability was 0.12 under five-fold cross-validation, and accuracy plateaued near 0.2 by 110 training accessions with no gain. Training-population size was not the limiting factor. The panel is a diverse collection rather than a structured breeding population, and this is an early-stage program. The phenomic advantage is not confined to a correlation coefficient. It changes which lines a breeder advances. The two selection directions converged as intensity relaxed, because a wider net forgives ranking error: at 20% and 30% retention, advancing and removal were indistinguishable. Handheld NIRS is therefore best deployed as an early-stage screen, advancing or removing materials cheaply and reserving Dumas reference chemistry for elite-tier confirmation, following the breeding logic that cheap predictors raise selection intensity early while expensive phenotyping is applied to a smaller advanced set (Heffner et al., 2009; Crossa et al., 2017; Bernardo, 2020).

### Laying the foundation for white lupin protein breeding

This study is an early step toward the goal of establishing and deploying a protein-breeding pipeline within Auburn University’s cover-crop and forage breeding program. Across four winter seasons, more than 400 USDA-NPGS accessions and Auburn breeding lines have been evaluated, and advanced yield trials have begun across Alabama environments. Anthracnose has not been a major constraint at our sites to date, but it remains an important breeding target in WL and has limited crop improvement and production in other environments (Schwertfirm et al., 2024; Annicchiarico et al., 2025).

Once a calibration is established, the phenomics platform delivers four practical benefits: (i) it reduces cost, recovering full calibration accuracy and heritability from roughly 40 of 125 Dumas assays, which run approximately $8 per sample including logistics (University of Missouri Agricultural Experiment Station Chemical Laboratories, 2026); (ii) it saves time, returning protein estimates for a full panel within a week, against the months of turnaround typical of outsourced Dumas combustion (approximately grinding 35 samples per day, Dumas proceeds at 20 samples per day); (iii) it simplifies phenotyping, requiring only a handheld scan of ground seed in-house rather than laboratory combustion; and (iv) it scales screening, allowing far more lines to be evaluated per season. Our results define how the two prediction tools are best used: handheld NIRS provides a rapid, low-cost screen for seed crude protein, while genomic selection remains positioned for parental choice and long-term gain as multi-season training populations accumulate. This reference-then-predict framework can extend beyond crude protein to amino-acid composition, seed oil, alkaloid content, and forage-quality traits within the program (Robert et al., 2022; DeSalvio et al., 2024; David et al., 2024).

### Limitations

These conclusions are specific to seed CP, the tested germplasm, and two seasons. The cross-year genetic correlation and cross-season genomic accuracy (r_g = 0.31; PA ≈ 0.12) indicate that GxY interaction, not marker quality, bounds temporal transfer; reliable forward prediction will require related, structured training material and multi-season training rather than denser genotyping.

## Conclusion

We compared the utility of phenomic and genomic prediction for seed CP in white lupin. Handheld NIRS predicted protein at screening-grade accuracy (R^2^ = 0.81; RPD = 2.34), transferred across years, and reached full calibration accuracy and heritability with 40 to 60 of 125 reference assays. Phenomic prediction and genomic prediction performance under identical validation (0.93 and 0.12), robustly across models and kernels; the weak genomic signal may reflect genotype-by-year interaction, germplasm structure, and limited within-panel relatedness. The result is a practical recommendation to use phenomics for rapid, low-cost screening, which will allow the application of genomic prediction for parent selection and increase accuracy, as program panel relatedness and multi-year training sets accumulate.

## Supporting information

Here

## Abbreviations

BLUE: best linear unbiased estimator
BLUP: best linear unbiased predictor
CP: crude protein
GRM: genomic relationship matrix
GS: genomic selection
GxY: genotype-by-year interaction
HPC: high-performance computing
MAE: mean absolute error
NIRS: near-infrared spectroscopy
PA: predictive ability
PLSR: partial least squares regression
PRM: phenomic relationship matrix
PS: phenomic selection
RKHS: reproducing kernel Hilbert space
RMSEP: root mean square error of prediction
RPD: ratio of performance to deviation
SNV: standard normal variate.

## Acknowledgement

This work was supported by the Southern Sustainable Agriculture Research and Education (SARE) program under grant GS24-315. The Alabama Wheat & Feed Grains Commission. We thank the Auburn University E.V. Smith Research Station and Plant Breeding Unit staff for field support. Dr. Sam Rochell, Denise Sanders, Julie Ross Datuin and Matias Machado of the Auburn University Department of Poultry Science for assistance with reference protein analysis. The USDA National Plant Germplasm System for providing WL accessions. Genotyping was performed using the KHUFU low-pass whole-genome sequencing pipeline (HudsonAlpha Institute for Biotechnology), Clevenger Lab and Sharon Riekhav et al. Computational analyses were performed on the Auburn University Easley High-Performance Computing Cluster.

## Data Availability

Low-pass WGS data for the white lupin panel are publicly available in the NCBI Sequence Read Archive under BioProject accession PRJNA1505808, titled “Genomic diversity and population genomics resource for white lupin breeding.

## Conflict of interest

The authors declare no conflict of interest.

## Supplemental Material

Seven tables support the main-text analyses. S1: predictive ability and RMSE for all seven models under three targets. S2: NIRS calibration performance (R^2^, RMSE, MAE, bias, RPD) for PLSR and three benchmarks, within-panel and cross-season. S3: genomic-phenomic kernel integration (RKHS) under BLUP and BLUE targets. S4: germplasm-stratified genomic prediction (Auburn, USDA-NPGS). S5: cross-season genomic prediction, both directions. S6: linear versus Gaussian kernel-form sensitivity (BLUE target). S7: training-population-size curve. S8: selection concordance at three intensities.

## Author Contributions

Mark Philip Castillo: Conceptualization, Methodology, Software, Formal analysis, Investigation, Data curation, Visualization, Writing - original draft, Writing - review & editing. Oluwaseye Gideon Oyebode: Methodology, Funding acquisition, Conceptualization, Investigation, Writing - review & editing. Ainsley Lenahan: Investigation, Data curation. Anthony Orloski: Investigation, Data Curation. Marnin Wolfe: Conceptualization, Methodology, Supervision, Resources, Funding acquisition, Writing - review & editing.

## Notes

### Competing Interest Statement

The authors have declared no competing interest.

## References

Adunola, P., Tavares Flores, E., Riva-Souza, E. M., Ferrão, M. A. G., Senra, J. F. B., Comério, M., Espindula, M. C., Verdin Filho, A. C., Volpi, P. S., Fonseca, A. F. A., Ferrão, R. G., Munoz, P. R., & Ferrão, L. F. V. (2024). A comparison of genomic and phenomic selection methods for yield prediction in Coffea canephora. The Plant Phenome Journal, 7(1), e20109. 10.1002/ppj2.20109.

Annicchiarico, P., Osorio, C., Nazzicari, N., Ferrari, B., Barzaghi, S., Biazzi, E., Tava, A., Pecetti, L., Notario, T., Romani, M., & Crosta, M. (2025). Genetic variation and genome-enabled selection of white lupin for key seed quality traits. BMC Genomics, 26, 922. 10.1186/s12864-025-12048-0

Barnes, R. J., Dhanoa, M. S., & Lister, S. J. (1989). Standard normal variate transformation and de-trending of near-infrared diffuse reflectance spectra. Applied Spectroscopy, 43(5), 772–777. 10.1366/0003702894202201

Barrett, T., Dowle, M., Srinivasan, A., Gorecki, J., Chirico, M., Hocking, T., Schwendinger, B., & Krylov, I. (2026). data.table: Extension of data.frame (R package version 1.18.2.1) [Computer software]. 10.32614/CRAN.package.data.table

Bates, D., Mächler, M., Bolker, B., & Walker, S. (2015). Fitting linear mixed-effects models using lme4. Journal of Statistical Software, 67(1), 1–48. 10.18637/jss.v067.i01

Bernardo, R. (2020). Breeding for quantitative traits in plants (3rd ed.). Stemma Press.

Brault, C., Lazerges, J., Doligez, A., Thomas, M., Ecarnot, M., Roumet, P., Bertrand, Y., Berger, G., Pons, T., François, P., Le Cunff, L., This, P., & Segura, V. (2022). Interest of phenomic prediction as an alternative to genomic prediction in grapevine. Plant Methods, 18, 108. 10.1186/s13007-022-00940-9

Braum, S. M., & Helmke, P. A. (1995). White lupin utilizes soil phosphorus that is unavailable to soybean. Plant and Soil, 176, 95–100. 10.1007/BF00017679

Breiman, L. (2001). Random forests. Machine Learning, 45(1), 5–32. 10.1023/A:1010933404324

Burns, D. A., & Ciurczak, E. W. (Eds.). (2001). Handbook of near-infrared analysis (2nd ed.). CRC Press. 10.1201/9781003042204

Caffarelli, P., Gastelle, J., Hunt, A., & Henderson, R. (2025). Domestic grain and oilseed transportation to the Southeastern United States (summary). U.S. Department of Agriculture, Agricultural Marketing Service. 10.9752/TS470.06-2025.

Chang, C. C., Chow, C. C., Tellier, L. C. A. M., Vattikuti, S., Purcell, S. M., & Lee, J. J. (2015). Second-generation PLINK: Rising to the challenge of larger and richer datasets. GigaScience, 4, 7. 10.1186/s13742-015-0047-8

Crossa, J., Pérez-Rodríguez, P., Cuevas, J., Montesinos-López, O., Jarquín, D., de los Campos, G., Burgueño, J., González-Camacho, J. M., Pérez-Elizalde, S., Beyene, Y., Dreisigacker, S., Singh, R., Zhang, X., Gowda, M., Roorkiwal, M., Rutkoski, J., & Varshney, R. K. (2017). Genomic selection in plant breeding: Methods, models, and perspectives. Trends in Plant Science, 22(11), 961–975. 10.1016/j.tplants.2017.08.011

Cullis, B. R., Smith, A. B., & Coombes, N. E. (2006). On the design of early generation variety trials with correlated data. Journal of Agricultural, Biological, and Environmental Statistics, 11, 381–393. 10.1198/108571106X154443

Daba, S. D., Honigs, D., McGee, R. J., & Kiszonas, A. M. (2022). Prediction of protein concentration in pea (Pisum sativum L.) using near-infrared spectroscopy (NIRS) systems. Foods, 11(22), 3701. 10.3390/foods11223701

David, L. S., Nalle, C. L., Abdollahi, M. R., & Ravindran, V. (2024). Feeding value of lupins, field peas, faba beans and chickpeas for poultry: An overview. Animals, 14(4), 619. 10.3390/ani14040619

DeSalvio, A. J., Adak, A., Murray, S. C., Jarquín, D., Winans, N. D., Crozier, D., & Rooney, W. L. (2024). Near-infrared reflectance spectroscopy phenomic prediction can perform similarly to genomic prediction of maize agronomic traits across environments. The Plant Genome, 17, e20454. 10.1002/tpg2.20454

Decision Innovation Solutions. (2021). Alabama economic analysis of animal agriculture: 2011–2021. Soy Transportation Coalition / U.S. Soybean Meal Demand Assessment. https://soymeal.org/demand-documents/2021/Alabama-Economic-Analysis-of-Animal-Agriculture-2011-2021.pdf.

Drucker, H., Burges, C. J. C., Kaufman, L., Smola, A. J., & Vapnik, V. (1997). Support vector regression machines. In M. C. Mozer, M. I. Jordan, & T. Petsche (Eds.), Advances in neural information processing systems 9 (pp. 155–161). MIT Press.

Endelman, J. B. (2011). Ridge regression and other kernels for genomic selection with R package rrBLUP. The Plant Genome, 4(3), 250–255. 10.3835/plantgenome2011.08.0024

Feldmann, M. J., Wang, F., & Runcie, D. E. (2026). Philosophy of phenomic prediction and its incompatibility with causal inference. The Plant Phenome Journal, 9, e70091. 10.1002/ppj2.70091.

Ferrari, B., Barzaghi, S., & Annicchiarico, P. (2022). Development of NIRS calibrations for seed content of lipids and proteins in contrasting white lupin germplasm. In X. Chu, L. Guo, Y. Huang, & H. Yuan (Eds.), 4Sense the real change: Proceedings of the 20th International Conference on Near Infrared (pp. 132–136). Chemical Industry Press. 10.1007/978-981-19-4884-8_13.

Forni, S., Aguilar, I., & Misztal, I. (2011). Different genomic relationship matrices for single-step analysis using phenotypic, pedigree and genomic information. Genetics Selection Evolution, 43, 1. 10.1186/1297-9686-43-1

García-Gudiño, J., López-Parra, M., Hernández-García, F. I., Barraso, C., Izquierdo, M., Lozano, M. J., & Matías, J. (2024). Use of Lupinus albus as a local protein source in the production of high-quality Iberian pig products. Animals, 14(21), 3084. 10.3390/ani14213084

Gebremedhin, A., Li, Y., Shunmugam, A. S. K., Sudheesh, S., Valipour-Kahrood, H., Hayden, M. J., Rosewarne, G. M., & Kaur, S. (2024). Genomic selection for target traits in the Australian lentil breeding program. Frontiers in Plant Science, 14, 1284781. 10.3389/fpls.2023.1284781

Gianola, D., & van Kaam, J. B. C. H. M. (2008). Reproducing kernel Hilbert spaces regression methods for genomic assisted prediction of quantitative traits. Genetics, 178(4), 2289–2303. 10.1534/genetics.107.084285

Gresta, F., Oteri, M., Scordia, D., Costale, A., Armone, R., Meineri, G., & Chiofalo, B. (2023). White lupin (Lupinus albus L.), an alternative legume for animal feeding in the Mediterranean area. Agriculture, 13(2), 434. 10.3390/agriculture13020434

Hamungalu, O., Abdollahi, M. R., Morel, P. C. H., Liu, S., & Wester, T. J. (2026). Determination of chemical composition and metabolizable energy of chickpea, faba bean, field pea, lentil and lupin compared to soybean meal for broiler chickens. Poultry Science, 105(2), 106286. 10.1016/j.psj.2025.106286

Heffner, E. L., Sorrells, M. E., & Jannink, J.-L. (2009). Genomic selection for crop improvement. Crop Science, 49(1), 1–12. 10.2135/cropsci2008.08.0512

Helgi Library. (2023, November 26). Which country produces the most lupins? https://www.helgilibrary.com/charts/which-country-produces-the-most-lupins/

Holland, J. B., Nyquist, W. E., & Cervantes-Martínez, C. T. (2003). Estimating and interpreting heritability for plant breeding: An update. Plant Breeding Reviews, 22, 9–112. 10.1002/9780470650202.ch2

Hufnagel, B., Marques, A., Soriano, A., Marquès, L., Divol, F., Doumas, P., Sallet, E., Mancinotti, D., Carrere, S., Marande, W., Arribat, S., Keller, J., Huneau, C., Blein, T., Aimé, D., Laguerre, M., Taylor, J., Schubert, V., Nelson, M., … Péret, B. (2020). High-quality genome sequence of white lupin provides insight into soil exploration and seed quality. Nature Communications, 11, 492. 10.1038/s41467-019-14197-9

Kaur, S., Pembleton, L. W., Cogan, N. O. I., Savin, K. W., Leonforte, T., Paull, J., Materne, M., & Forster, J. W. (2012). Transcriptome sequencing of field pea and faba bean for discovery and validation of SSR genetic markers. BMC Genomics, 13, 104. 10.1186/1471-2164-13-104

Korani, W., O’Connor, D., Chu, Y., Chavarro, C., Ballen, C., Guo, B., Ozias-Akins, P., Wright, G., & Clevenger, J. (2021). De novo QTL-seq identifies loci linked to blanchability in peanut (Arachis hypogaea) and refines previously identified QTL with low coverage sequence. Agronomy, 11(11), 2201. 10.3390/agronomy11112201.

Krause, M. R., González-Pérez, L., Crossa, J., Pérez-Rodríguez, P., Montesinos-López, O., Singh, R. P., Dreisigacker, S., Poland, J., Rutkoski, J., Sorrells, M., Gore, M. A., & Mondal, S. (2019). Hyperspectral reflectance-derived relationship matrices for genomic prediction of grain yield in wheat. G3: Genes, Genomes, Genetics, 9(4), 1231–1247. 10.1534/g3.118.200856

Lane, H. M., Murray, S. C., Montesinos-López, O. A., Montesinos-López, A., Crossa, J., Rooney, D. K., Barrero-Farfan, I. D., De La Fuente, G. N., Morgan, C. L. S., & Rooney, W. L. (2020). Phenomic selection and prediction of maize grain yield from near-infrared reflectance spectroscopy of kernels. The Plant Phenome Journal, 3(1), e20002. 10.1002/ppj2.20002

Laudadio, V., & Tufarelli, V. (2011). Influence of substituting dietary soybean meal for dehulled-micronized lupin (Lupinus albus cv. Multitalia) on early phase laying hens production and egg quality. 4Livestock Science, 140(1-3), 184–188. 10.1016/j.livsci.2011.03.029

Liaw, A., & Wiener, M. (2002). Classification and regression by randomForest. R News, 2(3), 18–22. https://CRAN.R-project.org/doc/Rnews/.

Liland, K. H., Mevik, B.-H., & Wehrens, R. (2026). pls: Partial least squares and principal component regression (R package version 2. 9–0) [Computer software]. 10.32614/CRAN.package.pls.

Lucas, M. M., Stoddard, F. L., Annicchiarico, P., Frías, J., Martínez-Villaluenga, C., Sussmann, D., Duranti, M., Seger, A., Zander, P. M., & Pueyo, J. J. (2015). The future of lupin as a protein crop in Europe. Frontiers in Plant Science, 6, 705. 10.3389/fpls.2015.00705

Mæhre, H. K., Dalheim, L., Edvinsen, G. K., Elvevoll, E. O., & Jensen, I.-J. (2018). Protein determination-Method matters. Foods, 7(1), 5. 10.3390/foods7010005

Meuwissen, T. H. E., Hayes, B. J., & Goddard, M. E. (2001). Prediction of total genetic value using genome-wide dense marker maps. Genetics, 157(4), 1819–1829. 10.1093/genetics/157.4.1819

Nalle, C. L., Ravindran, V., & Ravindran, G. (2012). Nutritional value of white lupins (Lupinus albus) for broilers: Apparent metabolisable energy, apparent ileal amino acid digestibility and production performance. Animal, 6(4), 579–585. 10.1017/S1751731111001686

Noffsinger, S. L., & van Santen, E. (2005). Evaluation of Lupinus albus L. germplasm for the Southeastern USA. Crop Science, 45(5), 1941–1950. 10.2135/cropsci2004.0575

Picard, R. R., & Cook, R. D. (1984). Cross-Validation of Regression Models. Journal of the American Statistical Association, 79(387), 575–583. 10.1080/01621459.1984.10478083.

Pérez, P., & de los Campos, G. (2014). Genome-wide regression and prediction with the BGLR statistical package. Genetics, 198(2), 483–495. 10.1534/genetics.114.164442

Pérez-Elizalde, S., Cuevas, J., Pérez-Rodríguez, P., & Crossa, J. (2015). Selection of the bandwidth parameter in a Bayesian kernel regression model for genomic-enabled prediction. Journal of Agricultural, Biological, and Environmental Statistics, 20(4), 512–532. 10.1007/s13253-015-0229-y

Piepho, H.-P., & Möhring, J. (2007). Computing heritability and selection response from unbalanced plant breeding trials. Genetics, 177(3), 1881–1888. 10.1534/genetics.107.074229

Quiñones, M. A., Lucas, M. M., & Pueyo, J. J. (2022). Adaptive mechanisms make lupin a choice crop for acidic soils affected by aluminum toxicity. Frontiers in Plant Science, 12, 810692. 10.3389/fpls.2021.810692

R Core Team. (2024). R: A language and environment for statistical computing (Version 4.4.1) [Computer software]. R Foundation for Statistical Computing. https://www.R-project.org/

Rife, T. W., Courtney, C., Hershberger, J., Shaver, B., Gore, M. A., Neilsen, M., & Poland, J. A. (2021). Prospector: A mobile app for high-throughput NIRS phenotyping. arXiv. 10.48550/arXiv.2104.07071

Rincent, R., Charpentier, J.-P., Faivre-Rampant, P., Paux, E., Le Gouis, J., Bastien, C., & Segura, V. (2018). Phenomic selection is a low-cost and high-throughput method based on indirect predictions: Proof of concept on wheat and poplar. G3: Genes, Genomes, Genetics, 8(12), 3961–3972. 10.1534/g3.118.200760

Robert, P., Brault, C., Rincent, R., & Segura, V. (2022). Phenomic selection: A new and efficient alternative to genomic selection. In N. Ahmadi & J. Bartholomé (Eds.), Genomic prediction of complex traits: Methods and protocols (Methods in Molecular Biology, Vol. 2467, pp. 397–420). Humana. 10.1007/978-1-0716-2205-6_14

Savitzky, A., & Golay, M. J. E. (1964). Smoothing and differentiation of data by simplified least squares procedures. Analytical Chemistry, 36(8), 1627–1639. 10.1021/ac60214a047

Schwertfirm, G., Schneider, M., Haase, F., Riedel, C., Lazzaro, M., Ruge-Wehling, B., & Schweizer, G. (2024). Genome-wide association study revealed significant SNPs for anthracnose resistance, seed alkaloids and protein content in white lupin. Theoretical and Applied Genetics, 137, 155. 10.1007/s00122-024-04665-2

Stevens, A., & Ramirez-Lopez, L. (2026). An introduction to the prospectr package (R package version 0.2.9) [Computer software]. 10.32614/CRAN.package.prospectr

Surma, A., Książkiewicz, M., Bielski, W., Kozak, B., Galek, R., & Rychel-Bielska, S. (2025). Development and validation of PCR marker array for molecular selection towards spring, vernalization-independent and winter, vernalization-responsive ecotypes of white lupin (Lupinus albus L.). Scientific Reports, 15, 2659. 10.1038/s41598-025-86482-1

Technow, F., Bürger, A., & Melchinger, A. E. (2013). Genomic prediction of northern corn leaf blight resistance in maize with combined or separated training sets for heterotic groups. G3: Genes, Genomes, Genetics, 3(2), 197–203. 10.1534/g3.112.004630

University of Missouri Agricultural Experiment Station Chemical Laboratories. (n.d.). Proximate analyses. https://aescl.missouri.edu/Prox.html.

USDA National Agricultural Statistics Service. (2022). Alabama county estimates: Soybean 2020–2021. USDA-NASS Southern Region. https://www.nass.usda.gov/statistics_by_state/Alabama/Publications/County_Estimates/2022/ALSoybean2022.pdf.

U.S. Soybean Export Council. (2026). Conversion table. https://ussec.org/buyer-tools/conversion-table/

VanRaden, P. M. (2008). Efficient methods to compute genomic predictions. Journal of Dairy Science, 91(11), 4414–4423. 10.3168/jds.2007-0980

Warsame, A. O., O’Sullivan, D. M., & Tosi, P. (2018). Seed storage proteins of faba bean (Vicia faba L.): Current status and prospects for genetic improvement. Journal of Agricultural and Food Chemistry, 66(48), 12617–12626. 10.1021/acs.jafc.8b04992

Wang, F., Feldmann, M. J., & Runcie, D. E. (2025). Do not benchmark phenomic prediction against genomic prediction accuracy. The Plant Phenome Journal, 8, e70029. 10.1002/ppj2.70029

Weiß, T. M., Zhu, X., Leiser, W. L., Li, D., Liu, W., Schipprack, W., Melchinger, A. E., Hahn, V., & Würschum, T. (2022). Unraveling the potential of phenomic selection within and among diverse breeding material of maize (Zea mays L.). G3: Genes, Genomes, Genetics, 12(3), jkab445. 10.1093/g3journal/jkab445

Wickham, H., François, R., Henry, L., Müller, K., & Vaughan, D. (2026). dplyr: A grammar of data manipulation (R package version 1.2.1) [Computer software]. 10.32614/CRAN.package.dplyr

Williams, P. (2014). The RPD statistic: A tutorial note. NIR News, 25(1), 22–26. 10.1255/nirn.1419

Wold, S., Sjöström, M., & Eriksson, L. (2001). PLS-regression: A basic tool of chemometrics. Chemometrics and Intelligent Laboratory Systems, 58(2), 109–130. 10.1016/S0169-7439(01)00155-1

Zapletal, D., Kudělková, L., Šimek, V., Jakešová, P., Macháček, M., Straková, E., & Suchý, P. (2017). Haematological indicators in hybrid mallard ducks (Anas platyrhynchos) with regard to the use of meal from whole white lupin seeds in their diet. Acta Veterinaria Brno, 86(3), 309–315. 10.2754/avb201786030309

Zhu, X., Leiser, W. L., Hahn, V., & Würschum, T. (2021). Phenomic selection is competitive with genomic selection for breeding of complex traits. The Plant Phenome Journal, 4(1), e20027. 10.1002/ppj2.20027

Zou, H., & Hastie, T. (2005). Regularization and variable selection via the elastic net. Journal of the Royal Statistical Society: Series B (Statistical Methodology), 67(2), 301–320. 10.1111/j.1467-9868.2005.00503.x

