## Supplementary material for "Near-infrared phenomic and genomic prediction for seed protein in winter legume white lupin (*Lupinus albus* L.): A utility comparison": Here

**Supplemental Tables**

**Supplementary Table S1**. Robustness of prediction across targets: BLUP and BLUE (7 models, 5-fold CV x 50 reps, n = 204).

| Source | Model | BLUE Acc | BLUE RMSE | BLUP Acc | BLUP RMSE |
|---|---|---|---|---|---|
| Markers (genomic) | BLUP | 0.12 | 2.132 | 0.14 | 0.846 |
| Markers (genomic) | BayesA | 0.104 | 2.163 | 0.136 | 0.855 |
| Markers (genomic) | BayesB | 0.107 | 2.156 | 0.138 | 0.853 |
| Markers (genomic) | PLSR | 0.110 | 2.287 | 0.157 | 0.889 |
| Markers (genomic) | Random Forest | 0.094 | 2.162 | 0.128 | 0.854 |
| Markers (genomic) | SVM linear | 0.115 | 2.263 | 0.162 | 0.881 |
| Markers (genomic) | SVM radial | 0.165 | 2.132 | 0.192 | 0.843 |
| Spectra (phenomic) | BLUP | 0.930 | 0.785 | 0.892 | 0.385 |
| Spectra (phenomic) | BayesA | 0.919 | 0.844 | 0.884 | 0.398 |
| Spectra (phenomic) | PLSR | 0.917 | 0.860 | 0.877 | 0.416 |
| Spectra (phenomic) | Random Forest | 0.930 | 0.806 | 0.901 | 0.376 |
| Spectra (phenomic) | SVM linear | 0.914 | 0.876 | 0.878 | 0.431 |
| Spectra (phenomic) | SVM radial | 0.876 | 1.069 | 0.877 | 0.417 |

**Supplementary Table S2.** NIRS calibration performance for seed crude protein (PLSR selected; benchmarks under identical preprocessing).

| Validation scheme | Model | n (test) | R2 | RMSE (% DM) | MAE (% DM) | Bias (% DM) | RPD |
|---|---|---|---|---|---|---|---|
| 80/20 (Within panel) | **PLSR** | 24 | 0.809 | 1.06 | 0.97 | -0.16 | 2.34 |
| 80/20 (Within panel) | Random forest | 24 | 0.787 | 1.12 | 0.97 | -0.10 | 2.22 |
| 80/20 (Within panel) | Elastic net | 24 | 0.779 | 1.14 | 1.00 | -0.06 | 2.17 |
| 80/20 (Within panel) | SVR radial | 24 | 0.741 | 1.24 | 1.07 | -0.19 | 2.01 |
| Cross-season (2024 to 2023) | Elastic net | 13 | 0.864 | 1.12 | 0.86 | -0.60 | 2.82 |
| Cross-season (2024 to 2023) | Random forest | 13 | 0.856 | 1.15 | 0.89 | -0.47 | 2.75 |
| Cross-season (2024 to 2023) | **PLSR** | 13 | 0.837 | 1.23 | 0.97 | -0.53 | 2.58 |
| Cross-season (2024 to 2023) | SVR radial | 13 | 0.827 | 1.26 | 1.06 | -0.67 | 2.51 |

R2, coefficient of determination on the test set; RMSE, root mean square error; MAE, mean absolute error; RPD, ratio of performance to deviation (SD of reference values / RMSE). PLSR (highlighted) was the calibration used for phenomic prediction; it is shown alongside alternative learners for reference. RPD values above 2.0 indicate calibrations suitable for screening, and values approaching 2.5 approach quantitative reliability. Reference protein was determined by Dumas combustion.

**Supplementary Table S3.** Genomic and phenomic kernel integration (RKHS; n = 204; 20 repeats of five-fold cross-validation).

| **Model** | **Kernel(s)** | **Repeats** | **BLUE Accuracy** | **SD** | **BLUP Accuracy** | **SD** |
|---|---|---|---|---|---|---|
| GS | Genomic (GRM) | 20 | 0.090 | 0.029 | 0.128 | 0.027 |
| PS | Phenomic (PRM) | 20 | 0.932 | 0.003 | 0.892 | 0.004 |
| G+P | Both kernels | 20 | 0.932 | 0.003 | 0.894 | 0.004 |

All models were fitted by reproducing-kernel Hilbert space regression (RKHS; BGLR, 20 repeats) with identical cross-validation partitions. Adding the genomic kernel gave no improvement over the phenomic kernel alone under either target (BLUP: 0.894 versus 0.892; BLUE: 0.932 versus 0.932). The response is the accession crude-protein value estimated as a random-effect BLUP or a fixed-effect BLUE; both incorporate NIRS-predicted records, and a validation against Dumas-measured protein alone is reported separately.

**Supplementary Table S4.** Within-group panel (five-fold cross-validation, 50 repeats; n = 204; accession crude-protein)

| Germplasm | n | GS | GS SD |
|---|---|---|---|
| AU lines | 70 | -0.231 | 0.076 |
| USDA-NPGS | 134 | -0.175 | 0.053 |

**Supplementary Table S5** Cross-seasong genomic prediction: train one season, predict the other, both directions (7 models).

| Direction | Train | Test | Model | Class | PA Mean | SD |
|---|---|---|---|---|---|---|
| Backward | 2024 | 2023 | **rrBLUP** | Linear mixed model | **0.121** | 0.031 |
| Backward | 2024 | 2023 | BayesB | Bayesian linear | 0.103 | 0.031 |
| Backward | 2024 | 2023 | BayesA | Bayesian linear | 0.096 | 0.025 |
| Backward | 2024 | 2023 | Random forest | Nonlinear ML | 0.095 | 0.028 |
| Backward | 2024 | 2023 | SVM linear | Machine learning | 0.149 | 0.040 |
| Backward | 2024 | 2023 | PLSR | Linear | 0.146 | 0.041 |
| Backward | 2024 | 2023 | SVM radial | Machine learning | 0.141 | 0.034 |
| Forward | 2023 | 2024 | rrBLUP | Linear mixed model | 0.143 | 0.062 |
| Forward | 2023 | 2024 | BayesB | Bayesian linear | 0.139 | 0.036 |
| Forward | 2023 | 2024 | Random forest | Nonlinear ML | 0.130 | 0.047 |
| Forward | 2023 | 2024 | BayesA | Bayesian linear | 0.125 | 0.034 |
| Forward | 2023 | 2024 | SVM radial | Machine learning | 0.209 | 0.024 |
| Forward | 2023 | 2024 | PLSR | Linear | 0.112 | 0.050 |
| Forward | 2023 | 2024 | SVM linear | Machine learning | 0.109 | 0.051 |

**Supplementary Table S6.** Kernel-form sensitivity (n = 204 BLUE target, 5-fold CV).

| Source | Linear PA | Gaussian PA |
|---|---|---|
| Genomic (markers) | 0.094 | 0.081 |
| Phenomic (spectra) | 0.921 | 0.913 |

**Supplementary Table S7 -** Training-population-size curve

| Training accessions | GS accuracy | SE |
|---|---|---|
| 30 | 0.088 | 0.036 |
| 50 | 0.112 | 0.036 |
| 70 | 0.179 | 0.026 |
| 90 | 0.195 | 0.023 |
| 110 | **0.197** | 0.024 |
| 130 | 0.192 | 0.024 |
| 150 | 0.195 | 0.024 |
| 164 (max) | 0.192 | 0.025 |

Sensitivity to training-population size was assessed by holding out a fixed test set of 40 accessions and drawing training sets of 30, 50, 70, 90, 110, 130, 150, and 164 accessions at random without replacement from the remaining 164. Each training size was sampled 30 times. Kernel models were fitted with mixed.solve on the training accessions and used to predict the held-out set; accuracy was the Pearson correlation between predicted and observed CP, averaged over the 30 samples. The maximum training size of 164 is the panel size (n = 204) minus the fixed test set.

**Supplementary Table S8.** Selection concordance in phenomic and genomic prediction (n = 204; 50 repeats of five-fold cross-validation). Both directions select the same number of accessions at each intensity, so the chance expectation is identical.

**Table S8a**

| Method | Direction | Intensity | Coincidence | SD | Efficiency |
|---|---|---|---|---|---|
| PS | advancing | 10% | 0.674 | 0.039 | 6.74 |
| PS | advancing | 20% | 0.749 | 0.019 | 3.74 |
| PS | advancing | 30% | 0.834 | 0.015 | 2.78 |
| PS | removing | 10% | 0.499 | 0.048 | 4.99 |
| PS | removing | 20% | 0.753 | 0.019 | 3.76 |
| PS | removing | 30% | 0.829 | 0.016 | 2.76 |

**Table S8b. (Advance versus cull)**

| $\pi$ | advance | cull | difference | meaningful |
|---|---|---|---|---|
| 10% | 0.674 | 0.499 | +0.175 | yes |
| 20% | 0.749 | 0.753 | −0.004 | no |
| 30% | 0.834 | 0.829 | +0.005 | no |

Integration, matched solver: Δ = +0.002.
